# Development and assessment of tailored illustrations to enhance community understandings of genetics topics

**DOI:** 10.64898/2026.03.17.711941

**Authors:** Audrey M. Arner, Tobias C. McCabe, Amanda Seyler, Siti Nurani Zamri, Tan Bee Ting A/P Tan Boon Huat, Kar Lye Tam, Patriciah Kinyua, Echwa John, Sospeter Ngoci Njeru, Yvonne A.L. Lim, Michael Gurven, Colin Nicholas, Julien Ayroles, Vivek V. Venkataraman, Thomas S. Kraft, Ian J. Wallace, Amanda J. Lea

**Affiliations:** Department of Biological Sciences, Vanderbilt University, Nashville, TN, USA, 37232; Department of Anthropology and Archaeology, University of Calgary, Calgary, Alberta, Canada; Department of Parasitology, Universiti Malaya, Kuala Lumpur, Malaysia 50603; Turkana Health and Genomics Project, Centre for Community Driven Research - Kenya Medical Research Institute, Nairobi, Kenya 54840-00200; Center for Community Driven Research, Kenya Medical Research Institute, Nairobi, Kenya 54840-00200; Department of Anthropology, University of California Santa Barbara, Santa Barbara, USA 93106; Center for Orang Asli Concerns, Subang Jaya, Malaysia 45790; Department of Integrative Biology, University of California Berkeley, Berkeley, California, USA; Department of Anthropology, University of Utah, Salt Lake City, Utah, USA; Department of Anthropology, University of New Mexico, Albuquerque, New Mexico, USA

## Abstract

**Objectives:** Effective communication about genetics concepts is essential for collaborative anthropological genetics research. However, communication can be challenging because many ideas are abstract and may be especially unfamiliar to communities with limited access to formal education. Indeed, there are no widely adopted models for communicating such information, nor a clear understanding of the social factors that may shape participant engagement. Here, we conducted a qualitative and quantitative, community-driven study to understand how illustrations can be useful to support concept sharing with two Indigenous groups—the Orang Asli of Peninsular Malaysia and the Turkana of Kenya.

**Methods:** We used a two phase approach to create and evaluate how illustrations can bolster communication about genetics concepts. First, we created images illustrating answers to frequently asked questions about genetics, iteratively updating the illustrations based on participant feedback. Second, we conducted 92 interviews to evaluate the finalized illustrations’ effectiveness. Finally, we analyzed the interview data using thematic analyses, multivariable modeling, and multiple correspondence analyses to identify patterns in participant understanding and feedback, including age, sex, market integration, and schooling.

**Results:** Participants reported high interest in genetics research (92%) and broadly positive perceptions of the illustrations. Familiar, locally-grounded imagery was preferred and associated with greater perceived clarity, while more technical illustrations were more frequently reported as confusing. Quantitative analyses showed strong internal consistency across measures of engagement and understanding, with modest variation by degree of market-integration, schooling, and sex.

**Discussion:** Our findings demonstrate that community-specific visualizations, co-developed through iterative feedback, can effectively support engagement with genetics research in participant communities.

## Introduction

Integrating genetics and genomics into biological and biomedical research has advanced our understanding of human evolution, disease, and phenotypic variation. However, such insights have not been equally spread across populations and have focused primarily on individuals of European ancestry living in high-income countries [1–4]. The importance of diverse sampling is widely recognized as crucial for robust understanding of both evolutionary processes and the genetic architecture of complex traits and diseases [5,6], and is essential to both address health disparities and maximize the reach of benefits from downstream discoveries [7]. This understanding has led to recent initiatives expanding genomic research in undersampled populations, for example, work from the H3Africa Consortium [8], Uganda Genome Resource [9], and BioBank Japan [10]. In contrast to large-scale initiatives, many anthropologists have built both health- and evolutionary-focused genomics studies around long-term relationships with individual Indigenous communities to promote communication, trust, and transparency [11–13]. Nevertheless, Indigenous populations around the world continue to remain some of the most underrepresented groups in any subfield of genomic research [3,4,14].

Indigenous populations are underrepresented in genetics research due to a complex interplay of factors, including historical practices of extraction and exploitation among both the biomedical and anthropological fields, with particularly high visibility cases in the United States [15,16], New Zealand [17], and southern Africa [18]. Thus, a lack of transparency about research goals, limited community engagement, misuse of samples, and failure to address community priorities have hampered participation in past genetic studies and continues to make many communities wary of participation today [19]. Within this context, several Indigenous communities and scholars have published strategies and recommendations for best practices, as well as explicit rules of engagement [20–23]. For example, in response to concerns about lack of informed consent, use of culturally-inappropriate language, and inadequate ethics review of research using their genetic data [18], the San people in southern Africa developed a code of research ethics. This code is centered around five broad principles — respect, honesty, justness and fairness, care, and process — and all research projects are reviewed against this code before they are approved [23]. At its core, this set of principles, as well as others that have been put forth [20,21], center around meaningful and continued engagement with participant communities during the research process. This engagement is predicated on effective communication of the science goals and processes to ensure communities understand the research being conducted, as well as its benefits and limitations.

Despite the acknowledged need to clearly communicate study goals, procedures, and results to participant communities, it can be difficult to discuss genetics topics, which are often complex, abstract, and tied to field-specific background knowledge and jargon [24]. For example, most concepts and processes in genetics are not visible to the naked eye, often making them less intuitive. Further, certain technical words that are used in English, such as “DNA”, “gene”, or “chromosome”, may lack direct equivalents in other languages [25]. Even some terms used to describe family relationships, such as “aunt” or “cousin”, may not have direct translations or may refer to different types of relationships in different cultures and languages [26]. Quantitative studies of genetics literacy further demonstrate that understanding of core genetics concepts varies widely across populations and can shift over time [27]. To combat these complexities, one way scientists have communicated genetics topics to both the general public and participant communities is through the use of images.

Images are one of the choice methods to convey genetics material because they can potentially be generalizable across cultures and languages, depict processes that may not be visible to the naked eye, and can be a starting point for concept sharing [28]. However, there are no guidelines or widely adopted models for how images should be developed or shared with participant communities. While a few examples have been published—specifically illustrations used for returning results [29,30] or as a supplement to informed consent [31]—these examples can be quite technical and include large amounts of field-specific terminology. Those that are more accessible rely on analogies, which can help provide visualization of abstract concepts by highlighting similarities to familiar phenomena [32]. For example, Arango-Isaza et al. used colored corn, which is an important crop with a long history of cultivation among the Mapuche communities they worked with, to explain genetic diversity and heritability [30]. Despite ongoing progress in this area, a remaining gap is that there is very little information in the literature about how images are developed, especially information about iterative engagement with communities and their requests and feedback. As a result, it also remains unclear whether engagement with and feedback on these materials are heterogeneous across audiences, and what community-specific or contextual factors are important to consider.

Here, we address these gaps by conducting a qualitative and quantitative, community-driven study to understand how illustrations can be useful to Indigenous communities interested in learning more about genetics, with the broader goal of improving engagement with genetics research and promoting collaborative approaches. To do so, we worked with two groups with which we have long-standing relationships through ongoing anthropological, genomic, and biological research studies: the Orang Asli, the Indigenous peoples of Peninsular Malaysia, and the Turkana, Indigenous pastoralists of northwest Kenya. We conducted this project in two phases. First, we created illustrations that address commonly asked questions about genetics from subsistence-level communities. The same set of images was initially piloted with both Orang Asli and Turkana community members across multiple rounds of fieldwork, and feedback was iteratively collected to update the illustrations. Although our initial goal was to develop broadly generalizable images appropriate for both Turkana and Orang Asli, this process revealed the importance of population-specific imagery and framing, leading us to prioritize community-tailored illustrations over generalizable illustrations (and thus focusing on Orang Asli; Turkana-specific images were not developed). Next, we presented the finalized, population-specific illustrations to Orang Asli communities and conducted interviews about the illustrations to evaluate their effectiveness. Finally, we explored interview responses to identify patterns in participants’ understanding and feedback, as well as the social and contextual factors shaping participants’ responses. Overall, this study responds to community-expressed interests in learning more about genetics and offers experiences for researchers seeking to engage in similar communication efforts.

## Methods

### Participant populations

#### Orang Asli

The Orang Asli are the Indigenous peoples of Peninsular Malaysia, comprising less than 1% (∼210,000 individuals) of the country’s population. They are typically divided into 19 distinct ethnolinguistic groups and three broad sub-groups, distinguished primarily by language, phenotype, and subsistence strategies [33]. Data for this study were collected from ten villages (Figure 1A) that are primarily situated in remote regions in the rainforest of Peninsular Malaysia and have historically experienced limited access to infrastructure, including formal education, transportation, and healthcare. The villages were predominantly occupied by members of the Batek, Jahai, Temiar, and Semai ethnolinguistic groups. The Batek and Jahai belong to the Negrito (Semang) sub-group, traditionally nomadic hunter-gatherers who speak Northern Aslian languages, while the Temiar and Semai belong to the Senoi sub-group, traditionally practicing swidden agriculture with a sedentary or semi-nomadic lifestyle and speaking Central Aslian languages [34].

**Figure 1:**
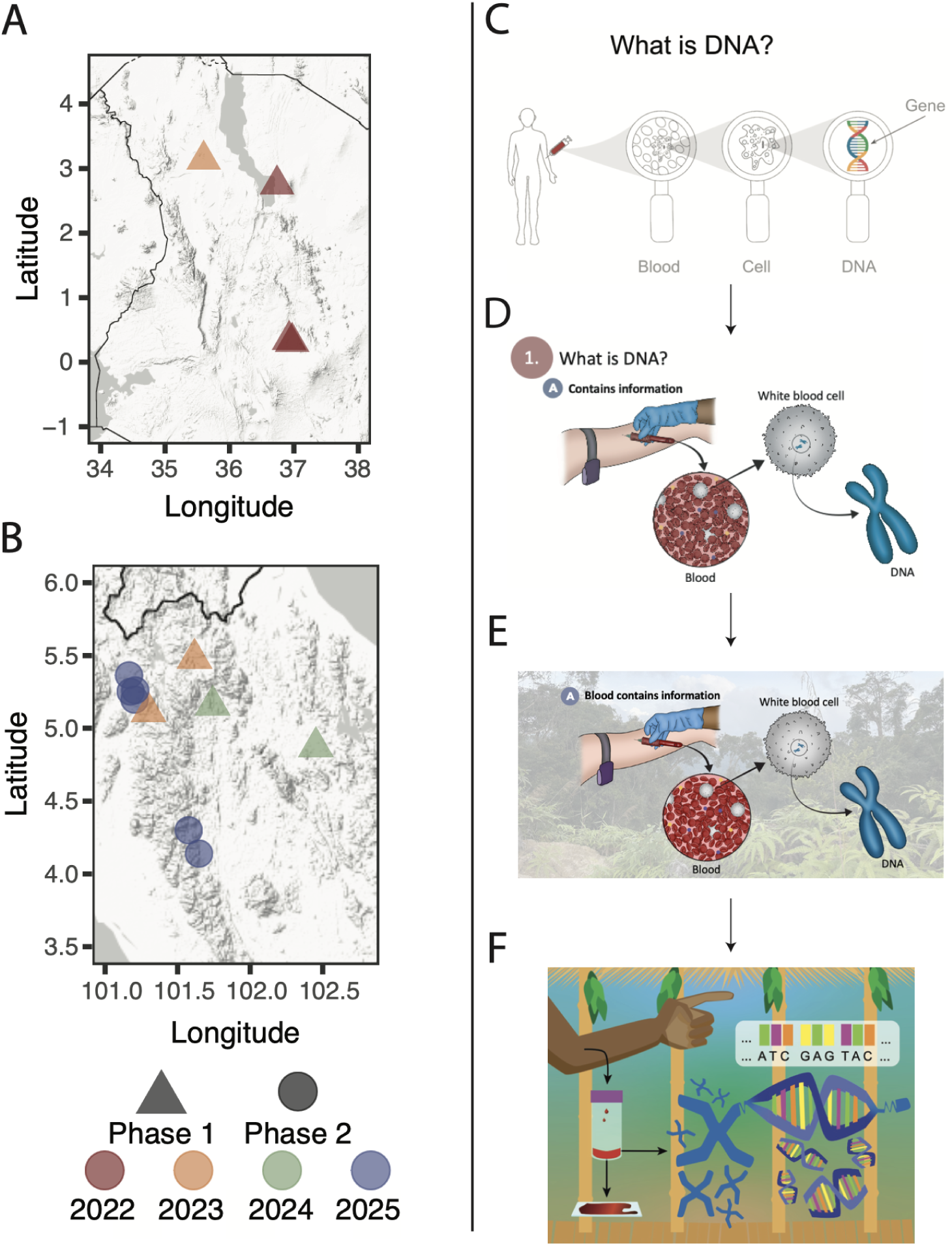
Study overview. (A-B) Map showing the locations where illustrations were piloted in Kenya and Malaysia respectively, with different colors representing the year the village was visited and shape representing the phase of the study. Phase 1 indicates piloting of images occurred at the marked location and Phase 2 indicates final presentations of the images and structured interviews at the marked location. (C-F) Development of the illustration answering the frequently asked question “What is DNA?”, where (C) is the first version presented and (F) is the final version presented. Although writing here is shown in English, it was written in Malay or Swahili in the presented images. When presented to participant audiences, Figure E would include photographs of Orang Asli.

Over the past 50 years, Malaysia’s rapid socioeconomic development has led to major lifestyle shifts for the Orang Asli, driven by two main forces: 1) the expansion of plantation agriculture and natural resource extraction has fragmented Orang Asli lands, and 2) government programs promoting assimilation into Malaysian society have shifted many Orang Asli away from traditional villages into organized resettlement schemes. The Orang Asli Health and Lifeways Project (OA HeLP) [35] is an international team of scientists, physicians, and Orang Asli advocates focused on how these lifestyle transitions are influencing health outcomes, using a range of questionnaire, anthropometric, biomarker, and genomic data types [36–39].

Data were collected from consenting adults ages 18 and older during trips conducted in partnership with OA HeLP, including 1) mobile clinics that visit communities to conduct research and provide free healthcare, or 2) follow-up trips to communicate results with study communities. However, participation in either concurrent or previous OA HeLP research and mobile clinics was not a requirement to participate in this study. Orang Asli communities included in this project were identified through existing relationships developed over many years of prior work by members of the OA HeLP team. The process of recruitment included two general steps. Permission to conduct the research was first sought from community leaders. This step was followed by the individual-level consent process, where the goals, research questions, and methods of this project were explained in detail after which formal, written consent was given.

#### Turkana

The Turkana are a nomadic pastoralist population living in the Turkana Basin in northwest Kenya. This study worked with Turkana individuals from four villages, two of which were remote and two of which were in more urbanized areas. Ongoing infrastructure construction and rapid economic development of Kenya has resulted in the growth of several urban centers in and near traditional Turkana lands, the expansion of small-scale markets, and an increased reliance on industry and agriculture. As a result, many Turkana no longer exclusively practice traditional pastoralism, instead relying on trade, small scale farming, and increasing participation in the market economy. In addition to socioeconomic changes happening within the Turkana region, many Turkana have moved to highly urbanized areas in central Kenya in the last several decades [40,41].

The Turkana Health and Genomics Project (THGP) is an international team of geneticists and biologists working to understand the health consequences of these transitions using integrated questionnaire, anthropometric, biomarker, and genomic data [37,42–44]. In partnership with THGP, informal interviews were collected from consenting adults ages 18 and older during either 1) mobile clinics that visit communities to conduct research and provide free healthcare, or 2) community engagement and outreach events. Participation in either concurrent or previous THGP research and mobile clinics was not a requirement to participate in this study. The study goals, research questions, and methods of this project were explained to participants in a language commonly used and understood within each study population by researchers before formal, written consent was given.

### Overview of project phases

The study protocol was reviewed by the BRANY Institutional Review Board (protocol no. 24-180-734) and was determined to be exempt. The OA HeLP was approved by the Medical Review and Ethics Committee of the Malaysian Ministry of Health (protocol ID: NMRR-20-2214-55565), the Malaysian Department of Orang Asli Development (permit ID: JAKOA.PP.30.052 JLD 21), and the Institutional Review Board of Vanderbilt University (protocol ID: 212175). The THGP was approved by Vanderbilt University (protocol ID: 00000162) and Kenya Medical Research Institute (KEMRI/SERU/CTMDR/119/4875).

Data collection occurred in two phases to develop illustrations and evaluate their effectiveness as a resource for communicating information about genetics research. In the first phase (Turkana-2022, Orang Asli-2023, Turkana-2023, Orang Asli-2024; Figure 2), we developed initial illustrations from a list of frequently asked questions (see below: Initial formulation of images), after which informal interviews were conducted to identify ways to improve the illustrations, using a mix of open-ended and multiple choice questions. In the second phase (Orang Asli-2025; Figure 2), we focused on updated, Orang Asli specific images and conducted structured interviews to assess participant response to the illustrations using yes/no, short-form open-ended, and ranking questions (see Supplementary Material). Finally, we analyzed the interview data collected in the second phase to identify patterns in participant feedback and responses.

**Figure 2:**
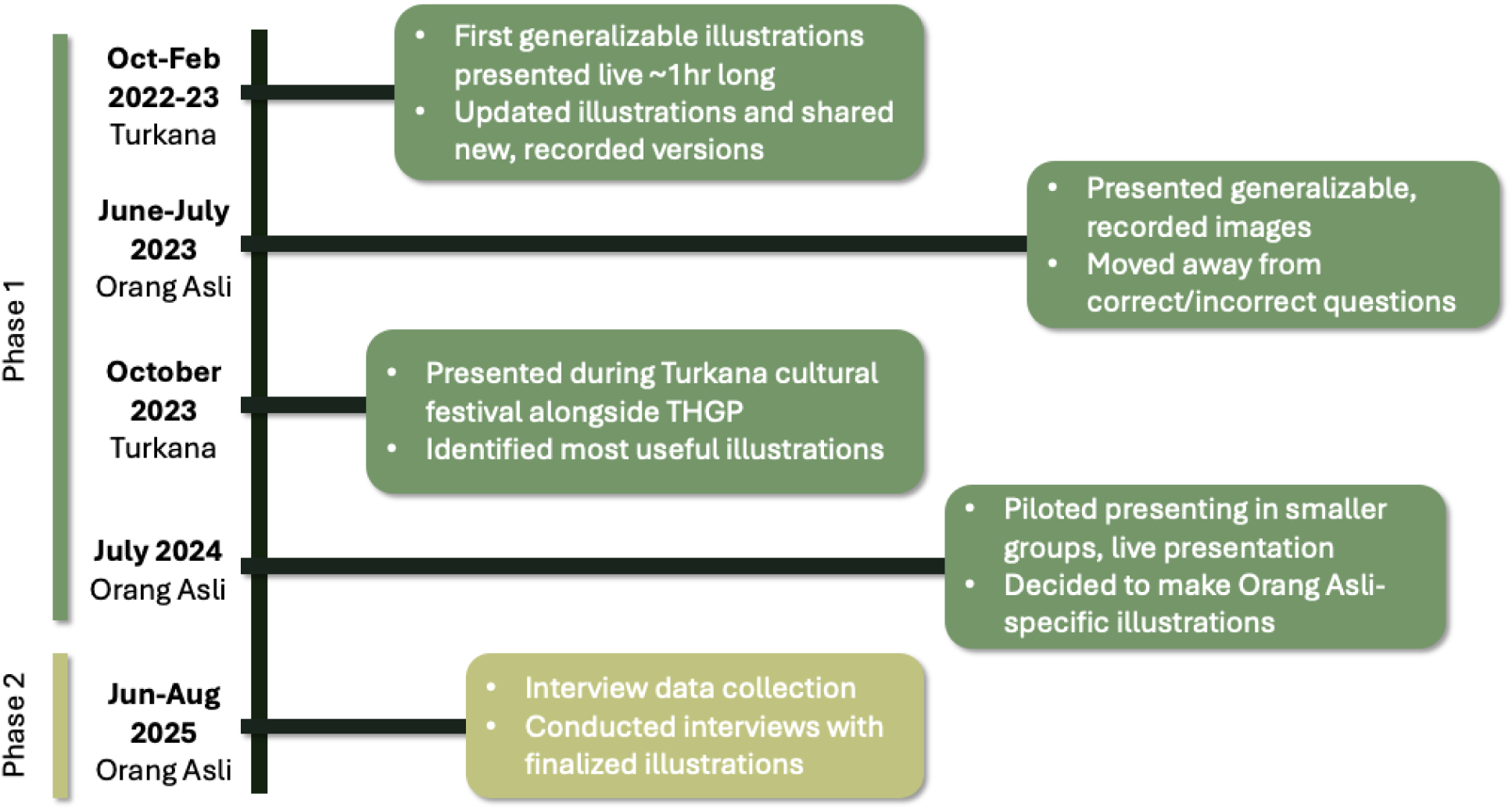
Timeline of illustration development, piloting, and evaluation across study phases. Key milestones in the iterative creation of genetics illustrations, including initial identification of frequently asked questions, piloting of generalizable illustrations, refinement of presentation formats, and the transition to Orang Asli–specific illustrations prior to formal interview-based evaluation.

### Initial formulation of images (Phase 1)

To design the initial illustrations, we began with a list of frequently asked questions raised by participants in a separate long-term anthropological and health research study with similar goals to OA HeLP and THGP (namely, the Tsimane Health and Life History Project [11]). This list of questions was the focus of project team and community discussions during follow-up results return trips for a recently published paper on Tsimane genetics [45]. We selected seven of these frequently asked questions to guide the development of the images: 1) What is DNA?, 2) Can DNA affect your health?, 3) What can scientists learn from DNA?, 4) Besides DNA, what else can you find from my blood?, 5) Why are scientists interested in markers of health in blood?, 6) What happens to my blood once you collect it?, and 7) Who has access to my DNA?

We created nine generalizable images to illustrate the above questions. To accommodate variable literacy levels, we minimized written text. All writing included on illustrations piloted in Turkana was written in Swahili, and writing on illustrations piloted with Orang Asli was written in Malay. Although Swahili and Malay are not the traditional languages within these groups, they function as regional lingua francas and are widely spoken in each area. We solicited feedback from community members and field assistants during two field seasons in Kenya (Turkana-2022, Turkana-2023) and two field seasons in Malaysia (Orang Asli-2023 and Orang Asli-2024). We used an iterative approach to incorporate community feedback, updating the images during each field season (Figure 1C). Ultimately, two versions of the illustrations were produced -- one generalizable version suited for multiple contexts (see https://github.com/audreyarner/genetic_illustrations), and one Orang Asli-specific version (https://github.com/tcmccabe/OrangAsliHealthIllustrations).

### Presentation of illustrations (Phase 1 and Phase 2)

In Phase 1, we piloted the dissemination of the illustrations using multiple formats--including live slide presentations, pre-recorded videos, tablet-based viewings, and printed discussion-based materials--to refine what was most useful to participants and could be reliably implemented, given that some locations do not have consistent access to electricity, wifi, or cell service. Similar to the illustrations themselves, the presentation of the illustrations was updated in an iterative manner, incorporating feedback to identify the most useful approach.

The illustrations were first disseminated at a Turkana community engagement event held in October 2022 (Turkana-2022). They were presented as a live talk, where a THGP research assistant presented the material in Swahili. Each illustration was pictured on a PowerPoint slide, and was shown via a projector. Later in the same field season (Turkana-2022), we presented a revised set of images into a 15-minute slide presentation that was narrated in Swahili and shown to individuals in three villages during mobile health clinics. The video format ensured a consistent presentation each time. The video format was also presented at the 2023 Turkana Cultural Festival, where THGP team members hosted a booth that highlighted their broader research activities and presented the illustrations (Turkana-2023).

We also presented the images in the 15-minute video format to Orang Asli communities (Orang Asli-2023). In this case, the video was pre-recorded with explanations in Malay and shown to community members in either large (∼20 people) community gatherings using a projector or small (2-5 people) group settings on a tablet (Figure 1B).

Based on feedback, we updated the presentation format such that OA HeLP research assistants led small discussion-based presentations of printed images, explaining each image to groups of one to six individuals (Orang Asli-2024). Each illustration was printed and laminated on A4 paper. Although the information conveyed varied slightly between sessions, this format allowed for interactive discussion and hands-on engagement with the materials. We engaged a broad cross-section of community members in these discussions, including Tok Batin (village headmen), teachers, community elders, and young adults who had recently completed secondary schooling. To ensure accessibility beyond digital settings and as a future resource, we also produced printed pamphlets featuring the same content (Orang Asli-2024).

In Phase 2 (Orang Asli-2025), the final illustrations were shown during a live presentation in a medium-large group setting to evaluate efficacy and gather participant feedback. Similar to earlier formats, this presentation used either PowerPoint slides (when electricity was available) or laminated, printed versions for the participants to view. These presentations were delivered live in Malay by an OA HeLP research assistant, which allowed for interactivity and interruptions if viewers had questions. The research assistant involved in Phase 2 did not have a background in genetics, but instead had discussed the image explanations with a researcher who had a background in the field to improve phrasing and understandability. The final presentations lasted approximately 12 minutes. The presentation was delivered in six Orang Asli communities, with between 20 and 70 community members attending each session. We again provided pamphlets featuring the same content to participants for future reference.

### Interviews (Phase 1 and Phase 2)

In Phase 1, we conducted brief, unstructured interviews during four field seasons (Turkana-2022, Orang Asli-2023, Turkana-2023, Orang Asli-2024) to assess how the illustrations, their presentation, and the interview itself could be improved to best meet community needs. Interviews were conducted in either Malay with Orang Asli participants or Swahili with Turkana participants, and individuals were free to answer whichever questions they wanted. Questions included open-ended items about what individuals liked most and least about the images, as well as gain of knowledge questions assessing understanding of some of the illustrated genetics concepts.

In Phase 2, we conducted structured, short format interviews with Orang Asli participants to collect both qualitative and quantitative data (Orang Asli-2025). These interviews focused on four main areas: demographic information, prior knowledge before viewing the presentation, opinions of the illustrations, and perceived knowledge empowerment (see Supplementary Information). Several questions were refined from those piloted during earlier field seasons. All interviews were conducted in Malay by local OA HeLP research assistants and lasted approximately 10-15 minutes. In total, 92 participants across six villages completed the structured interviews (SI Table 2).

### Thematic analysis of responses to open-ended questions (Phase 2)

To identify broad ideas underlying responses to the three open-ended interview questions, we conducted an iterative, inductive thematic analysis [46] of participant responses to each question separately. Specifically, two researchers (A.M.A. & A.S.) independently open-coded all responses to each question inductively using MAXQDA version 26 software to identify recurring concepts and patterns. A given response could have multiple phrases conveying different ideas. Therefore, the unit of analysis in coding was a phrase connected to an idea. Initial coding of the responses to each interview question was discussed, where researchers systematically reviewed code definitions and applications, with discrepancies resolved by consensus. Percent agreement was calculated to assess coding consistency between researchers, which ranged from 86% to 100% agreement (SI Table 3). After consensus coding was reached, researchers independently identified higher-order themes that emerged from the identified codes, followed by reflexive discussion to clarify and refine thematic structure and description for each theme. The most common themes were identified based on number of mentions (SI Figures 1-3).

### Statistical analysis of interview data (Phase 2)

First, we used binomial models to test whether the proportion of individuals responding affirmatively to each yes/no question differed from that expected by chance, fitting models separately for each question. We corrected for multiple hypothesis testing using a Benjamini-Hochberg false discovery rate [47]. We also calculated a pairwise Pearson correlation matrix to evaluate how responses to each question co-varied across individuals.

Second, we tested whether answers to the questions in our interviews could be composed into delineable axes of variation (e.g., whether groups of individuals tended to answer certain questions similarly). To do so, we used a multiple correspondence analysis (MCA), a type of exploratory factor analysis designed to reduce the dimensionality of categorical data [48]. Our MCA included the pool of the eight yes/no questions converted to Boolean format (true/false). Two individuals were removed due to one or more missing answers, resulting in a total of 90 individuals for this analysis. MCA was performed using the FactoMineR package in R with default parameters [49]. The first two dimensions were retained based on their relative inertia (35.3% and 18.9% respectively; see SI Figure 4) and interpretability.

We next tested whether individual coordinates on MCA dimensions 1 and 2 were shaped by any sociodemographic factors, namely sex, age, highest education level (coded as a linear variable with 0 representing no formal education, 1 representing some primary education, 2 representing some secondary education, and 3 representing some university education), and “urbanicity”. Here, we used a location-based “urbanicity score” that captures access to industrialized, market-based resources available across the community (e.g., access to electricity, sewage, formal education; see Supplementary Text for urbanicity score generation). This score was first proposed by Novak et al [50] and has previously been tested in Orang Asli [36]. We used linear models including sex, age, highest education level, and urbanicity score to predict MCA dimensions 1 and 2 in separate models [47]. We also ran follow-up models in which highest education level was coded as a binary variable of no formal education (coded as 0) versus any level of formal education (coded as 1).

Finally, we used linear models to analyze whether demographic and other factors impacted response to each yes/no question. For each question separately, we fit a binomial model in which response (yes/no) was predicted jointly by age, sex, highest level of schooling, or urbanicity score. We again corrected for multiple hypothesis testing using an FDR approach. Similar to the above, we ran follow-up models switching highest education level with a binary variable of no formal education versus any level of formal education. All analyses were performed using the R computing language and RStudio (version 4.2.1).

## Results

We developed a series of illustrations to address frequently asked questions about genetics. Both a broad, generalizable version and an Orang Asli-specific version of the illustrations are available on GitHub (https://github.com/audreyarner/genetic_illustrations, https://github.com/tcmccabe/OrangAsliHealthIllustrations) and are also accessible from the OA HeLP project website (https://www.orangaslihealth.org/). In the following text, we describe: Phase 1, which included the iterative development and refinement of the illustrations; and Phase 2, which included the presentation of the final version of illustrations and the qualitative and quantitative evaluations of their efficacy and drivers of engagement.

### Phase 1: Iterative development of illustrations depends on community feedback

Given the desirability of a generally-applicable genetics resource, our first round of pilot images depicted people, objects, and environments that were not specific to any geographic region (Figure 1C). For example, we used a simplified human outline without any identifiable phenotypes or characteristics to enhance relatability across contexts, consistent with genetics imagery used in prior publications [29,30]. A key theme in early feedback (Turkana-2022) was the desire for more realistic images. In response, we revised the illustrations to include greater visual detail and less abstract depictions of individuals, including a range of skin tones (Figure 1D). Feedback on the revised illustrations was generally positive; however, viewers found the mode of illustration dissemination (a video; Turkana-2023, Orang Asli 2023), to be too long and insufficiently interactive. Therefore, participants suggested the inclusion of more dynamic elements such as animation. Similar to feedback from Turkana, Orang Asli viewers suggested that the video was too long, with viewers reporting that this style of presentation was not interactive enough (Orang Asli-2023). Orang Asli feedback also provided new perspectives, highlighting a desire for the presentation to include imagery that was more locally relatable and specific to their lives.

Based on this community feedback, we prioritized shortening the presentation, making the presentation more interactive, and incorporating Orang Asli-specific elements into the illustrations for additional piloting (Figure 1E, Orang Asli-2024). To shorten the presentation we removed one of the images (which answered the question “What happens to my blood once you collect it”) that was the most repetitive. For each image, we included relevant pictures taken in Orang Asli communities, as well as added a rainforest background to each of the illustrations. The most common suggestion we received at this stage was to incorporate additional Orang Asli-specific examples and imagery, as well as examples of genetics principles that community members would be more familiar with. Although our original goal was to produce a generalizable resource suitable for multiple populations, this feedback prompted a pivot toward developing an Orang Asli-specific version as the primary set of illustrations.

Feedback also informed the development of our interview questions. During early pilot interviews (Turkana-2022, Orang Asli-2023), some community members noted that it felt like taking a test, which they remarked was not enjoyable. Because our goal was to assess how effective the illustrations were for knowledge empowerment rather than acquisition by Western standards, we removed items with definitive “right” or “wrong” answers. Additionally, we limited the number of open-ended questions included in the final interview, as we found that individuals had a difficult time putting some concepts into words without prompts. Overall, community feedback was consistently constructive, emphasizing appreciation for the visual and oral approach and the perceived increasing relevance of the illustrations to their own experiences.

### Phase 2: Final content and presentation focused on population-specific imagery

The finalized illustrations were structured around six frequently asked questions about genetics (Figure 3; SI Table 1; Orang Asli-2025). Guided by community recommendations, this Orang-Asli specific version incorporated recognizable local examples of genetics. For instance, hair texture -- which is highly variable within Orang Asli ethnolinguistic groups -- was used to illustrate heredity (Figure 3B), replacing height which shows less visible local variation. Additionally, we used durian, a popular fruit in Malaysia that has many easily-recognizable varieties varying in appearance, texture, and taste, to explain genetic diversity and the impact the environment can have on traits (Figure 3C). Because small-scale farming and close interaction with cultivated and wild plants are a part of daily life for many Orang Asli communities, this example leveraged shared experiential knowledge to make genetic variation more accessible. Similar locally grounded adjustments were made across all illustrations, including depicting people in traditional clothing and housing.

**Figure 3:**
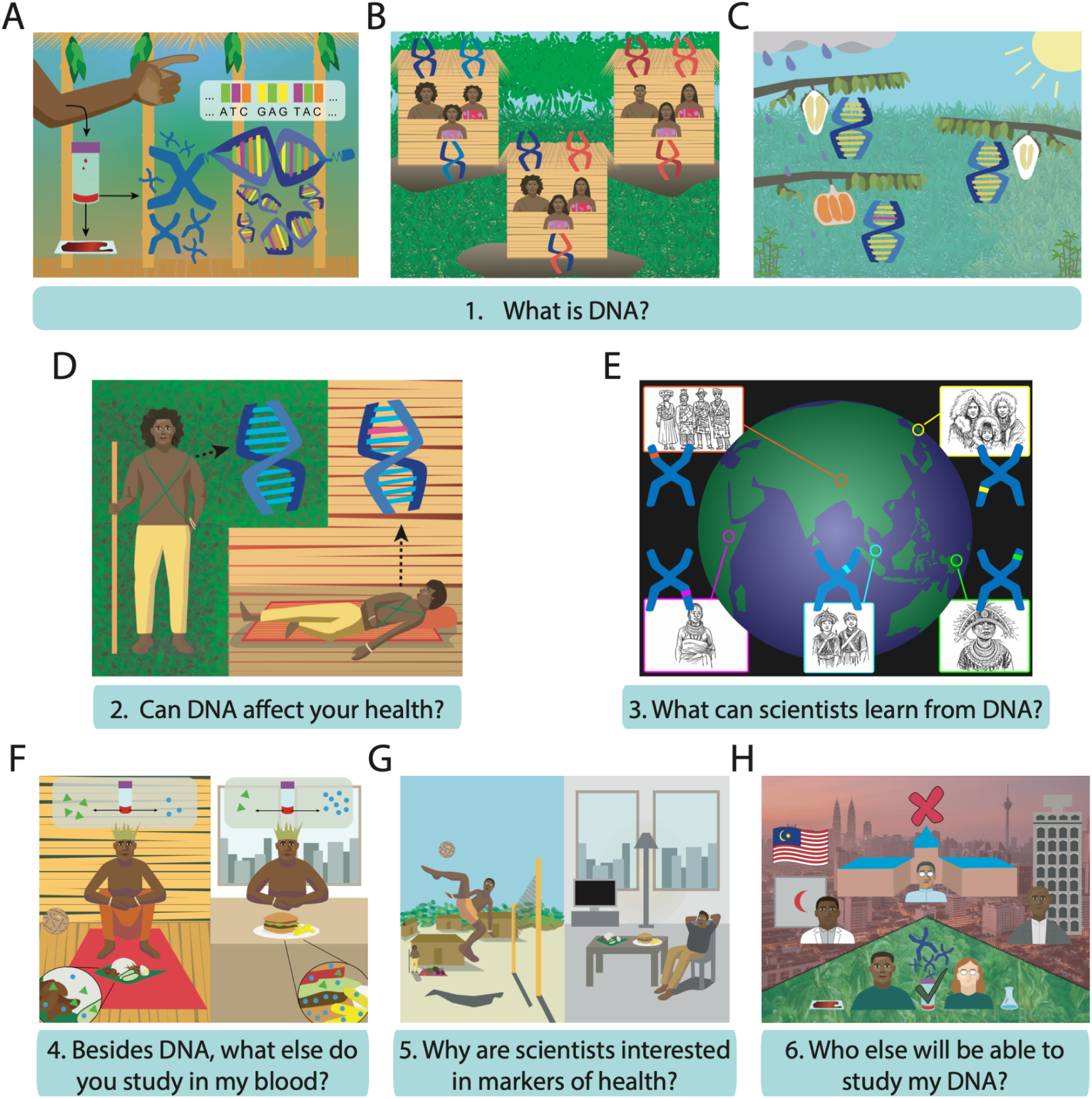
Final, Orang Asli-specific illustrations presented to communities. Below each image, we specify the frequently asked question being depicted. Throughout the text and in other figures, we refer to each image by their associated letter here (A-H). When presented, Illustration E included photographs of other Indigenous populations from the locations specified.

### Phase 2: Illustrations reported as useful by participants

To understand whether the illustrations were helpful in relaying genetics concepts, we conducted structured interviews with 92 Orang Asli individuals (Supplementary Table 2). In total, 85 participants (92%) reported wanting to know more about genetics research, and 44 participants (48%) reported they had believed prior to seeing the illustrations that there was health-related information in their blood. Participants answering affirmatively to the second question were asked a follow-up about the type of information they believed would be present. Nine individuals did not have a specific response. For those who responded, we used a thematic analysis to identify two major themes that developed from the data (SI Figure 1). First, participants described knowledge of measurable indicators coming from blood, often referencing specific biological markers or tests (e.g., “blood has sugar in it”). Second, participants expressed knowledge that blood can be used to assess medical status, reflecting broader health interpretations (e.g., “diseases and health”).

#### Qualitative analysis: Illustration preferences align with familiarity, while technical images are more confusing

We then asked questions about the illustrations to understand participants’ preferences and points of confusion. The greatest percentage of participants (38%) reported Illustration B as their favorite (Figure 3B). This image is one that Orang Asli would likely be the most familiar with, depicting the inheritance of hair texture -- a trait with observable variation among Orang Asli -- showing individuals residing with their family in traditional bamboo houses set in a rainforest environment. To formally assess why certain images were preferred, we conducted a thematic analysis of participants’ open-ended explanations for their favorite images (SI Figure 2). The most common theme (n=54 mentions) was preference for imagery related to identity, with participants frequently referencing recognizable environments and lived experiences, noting, for example, that the illustration “is the same as my daily activities, like playing takraw”. Participants also emphasized an interest in genetics concepts (n=31 mentions), explaining that they liked some images because they “liked knowing that everyone has their own DNA”. Smaller subsets of participants expressed preference for the visual and aesthetic components (e.g., “the picture is beautiful and elegant”; n=10 mentions), health-related imagery (e.g., “because the picture shows how you can be healthy”; n=25 mentions), and depictions that there are lifestyle-related health benefits (e.g., “healthy lifestyle of the village”; n=25 mentions).

We also asked which images, if any, were confusing to viewers (Figure 4B); 85% of participants reported at least one image as confusing (mean=1.8 confusing images). For example, the image most often reported as confusing (selected in 30% of all image responses, with participants able to select multiple images) was Illustration E, which depicts DNA variation in other Indigenous communities. This image was the most technical; however, it was retained in the final set of illustrations given feedback from Phase 1 to include information about other Indigenous populations around the world.

**Figure 4.**
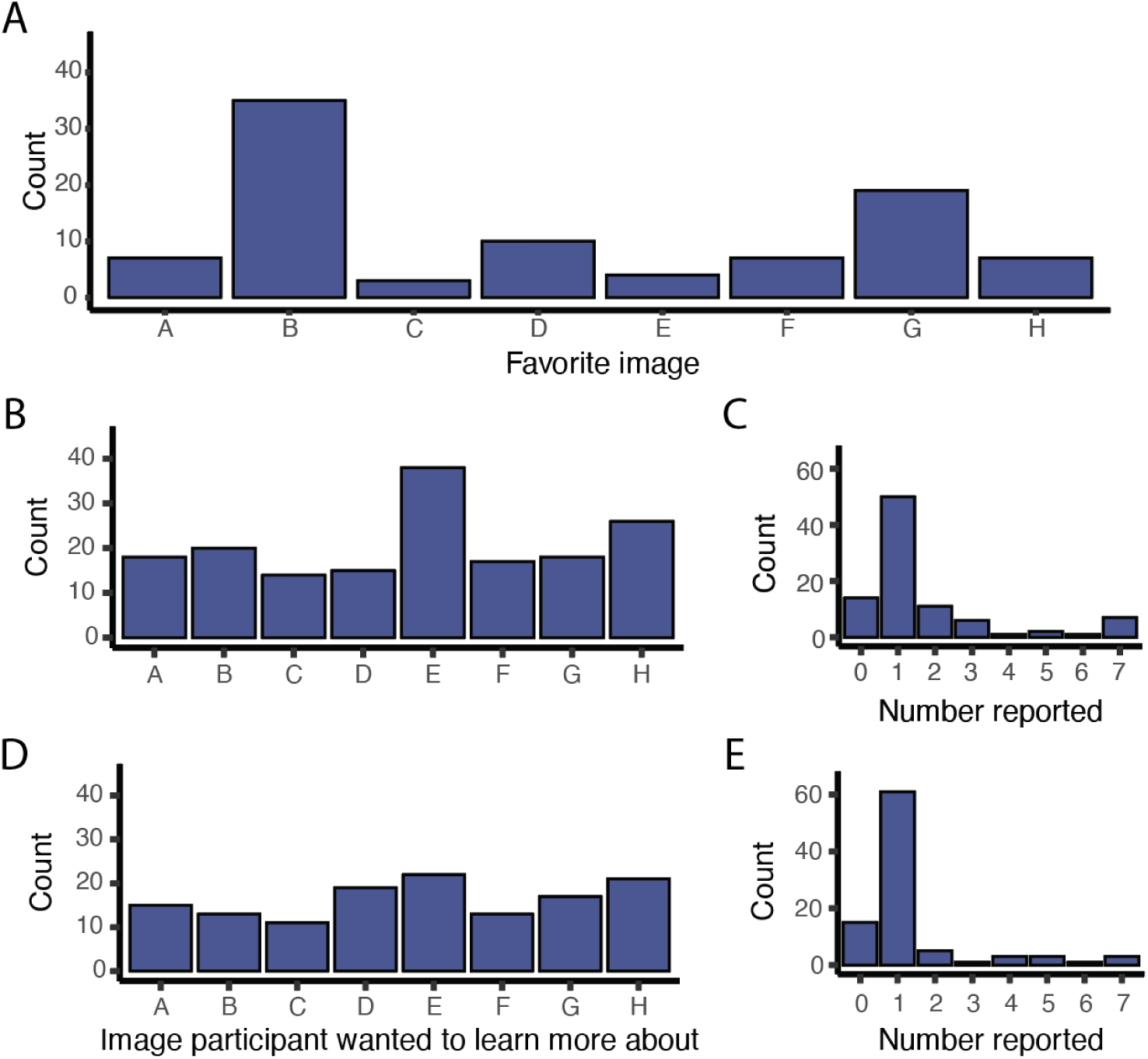
Participants’ preference and interests in genetics illustrations. Illustrations labeled according to Figure 3. (A) Barplot of participants’ favorite image. Each participant could select only one illustration as their favorite. (B) Barplot indicating which images were confusing. Participants could select multiple images (C) Barplot showing the number of images each participant reported as confusing. (D) Barplot indicating which images participants wanted to know more about. (E) Plot depicting the number of images each participant chose, with multiple illustrations able to be chosen.

In order to understand topics of continued interest, we asked what images, if any, participants would want to learn more about. Most participants reported wanting to learn more about at least one image (mean=1.4 images). Interest was fairly evenly distributed across the different illustrations (Figure 4C). As a follow up, we asked what general topics participants would like to have learned more about (SI Table 4). Most participants expressed interest in learning more about health and disease (49% of individuals) and relatedness (46% of individuals). A small number of participants selected “other”, primarily raising questions about blood type.

Finally, we asked participants what one thing they learned from the illustrations was. We again used a thematic analysis of short-form open-ended responses, which revealed four main themes (SI Figure 3). The most common theme reflected increased understanding of the role of DNA and blood in the body, with participants describing new awareness that blood contains biological information and that DNA influences bodily traits and health (n=73 mentions). For example, some participants noted “everyone has their own DNA” and “DNA changes can impact health”. A second theme involved factors contributing to health and well-being (n=32 mentions). Responses referenced learning, for example, that “blood has health information.” The third theme captured recognition of genetic variation among individuals and populations (n=21 mentions). Finally, a smaller but important theme reflected awareness of a knowledge gap, with participants identifying difficulty articulating a specific concept they learned or noting they wanted to learn more in the future (n=7 mentions).

#### Quantitative analysis: Genetics illustrations were broadly engaging and improved understanding, and engagement showed modest variation by education, sex, and urbanicity

We sought to understand the effectiveness of the illustrations for participants’ wants by asking eight yes/no questions assessing self-reported interest and knowledge gain. All questions were answered affirmatively more than expected by chance (FDR<0.05), indicating that participants found the illustrations engaging, understandable, and informative more so than not (Figure 6A, SI Table 5). We next assessed the correlation between answers (Figure 6B). Responses showed internal consistency, with measures of self-reported understanding and engagement positively correlated with one another (e.g., “I would view the images again” and “I would recommend to a friend”). Reporting that at least one of the illustrations was hard to understand was negatively correlated with nearly all other questions, particularly questions related to engagement and recommendation to others (mean Pearson r = -0.2).

**Figure 6.**
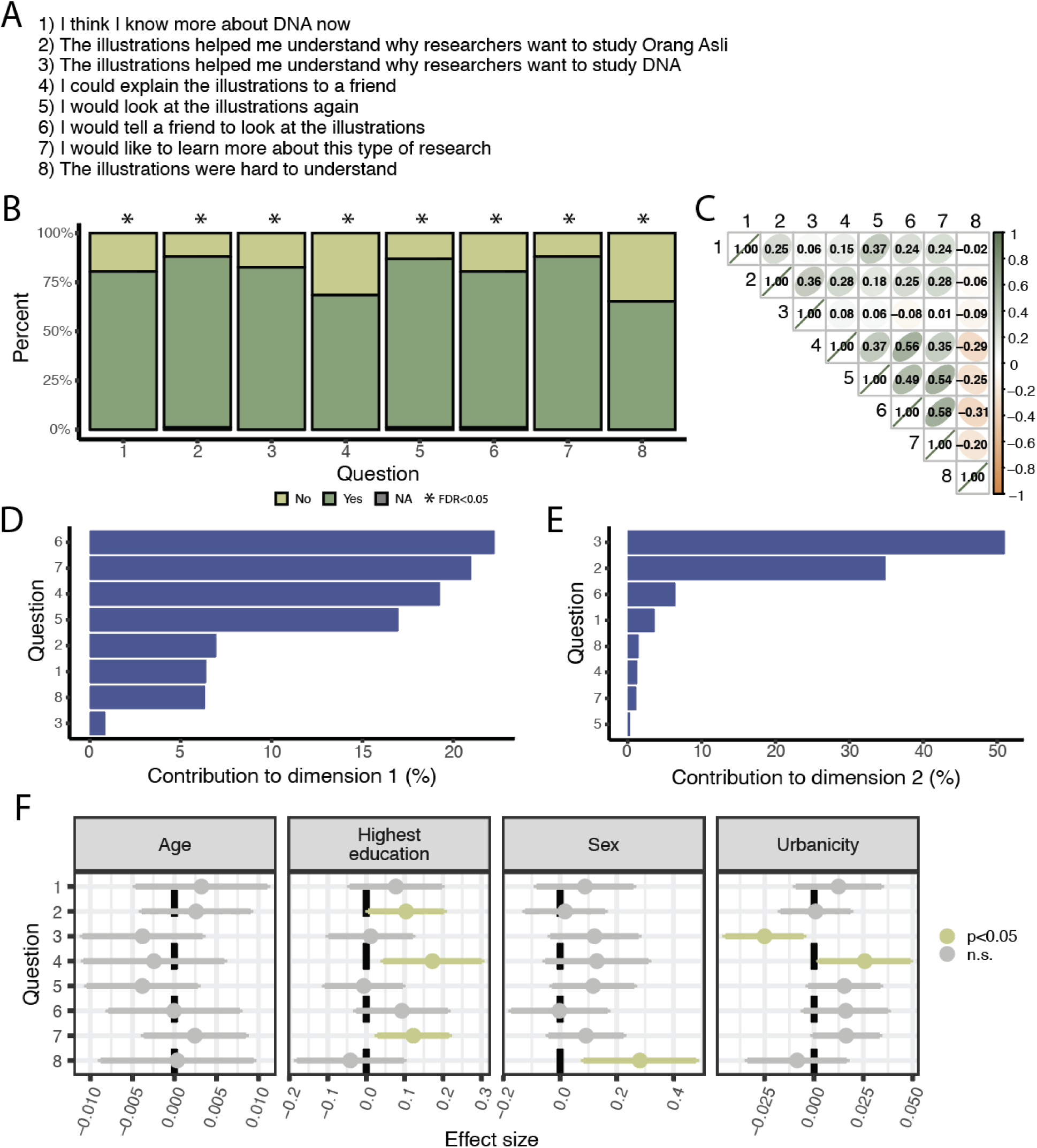
Self-reported satisfaction is high for community-specific genetics illustrations. (A) Questions participants were asked. Their number of 1 through 8 is repeated across panels of the figure. (B) Barplot showing the percentage of participants who answered “yes” vs “no” for each question used to determine effectiveness of illustrations. Binomial modeling was used to test whether each proportion was significantly different than 0. (C) Correlation between participant answers to each question. Numbers correspond to the questions in panel A. (D-E) Loadings of each question in MCA for dimensions 1 and 2 respectively. (F) Forest plot showing effect size and confidence interval for each demographic covariate of interest using binomial modeling. Color represents significance threshold.

To further explore correlations in participants’ responses, we performed a multiple correspondence analysis (MCA) using all eight yes/no questions. We determined that two dimensions accounted for the majority of the variance (dimension 1 proportion of variance: 35.3%, dimension 2 proportion of variance: 18.9%; SI Figure 1). The first dimension appeared to primarily reflect interest and engagement, with high loadings for items such as “I would recommend the illustrations to a friend” and “I want to learn more” (Figure 6C). The second dimension was driven by questions related to understanding and clarity, loading more strongly on questions such as “the illustrations helped me understand more about genetics” and “I understand why scientists would want to study DNA” (Figure 6D). We then modeled associations between individuals’ scores on the first two MCA dimensions and four predictors: age, sex, highest attained level of education, and urbanicity. Although none of the predictors remained significant after multiple hypothesis testing correction, dimension 2 was nominally associated with urbanicity score (p=0.042) (SI Table 6).

Finally, we modeled each individual interview question as a function of age, sex, highest attained level of education, and urbanicity (Figures 6E, SI Table 7). While no predictors remained significant after multiple hypothesis testing correction, we found that highest education level showed the most consistent associations, reaching a nominal P<0.05 for three questions. A higher level of education was associated with “yes” responses for all three of these questions (Figure 6F). Additionally, we found that individuals with lower urbanicity were more likely to report that the illustrations helped them understand why researchers want to study DNA (P=0.03; SI Table 7), while individuals with lower urbanicity were more likely to report that they could explain the illustrations to a friend (P=0.003; SI Table 7). Finally, men were more likely than women to report that some of the illustrations were hard to understand (P=0.01; SI Table 7). We found that results were very similar when using a binary of none vs any formal education (SI Table 8). Given the importance of specific histories, we ran additional models including ethnolinguistic group as a predictor. Together, these patterns suggest that prior exposure to formal education and biology concepts, which vary systematically with urbanicity, can influence how individuals interpret and engage with genetics communication materials.

## Discussion

Effective communication of genetics information is essential to ethical research partnerships [20,23]. However, genetics concepts are often abstract, technical, and challenging to convey across diverse linguistic and cultural contexts. Visualization strategies can bridge this gap by connecting intangible concepts to concrete, visible examples. While a few examples of illustrations [29–31] and videos [51,52] explaining genetics concepts to participant communities have been published, there is limited information about how these visuals are developed or received. Here, we used an iterative, community-based process to demonstrate that illustrations can serve as effective tools for communicating genetics research with Indigenous communities, but that engagement and comprehension are shaped by demographic and contextual factors.

First, we found that iterative, community-driven development was essential for producing illustrations that aligned with participant priorities. The initial images we created (Figure 1C) differed substantially from the final versions (Figure 1F), with multiple rounds of revision informed by community feedback. This process aligns with participatory research approaches that emphasize co-creation and responsiveness to users rather than one-directional knowledge transmission. Indeed, prior work in science communication and community-based participatory research has shown that iterative development improves relevance, trust, and engagement, particularly when communicating complex or sensitive topics [53,54]. Interestingly, this iterative process revealed that participant priorities did not always align with maximizing simplicity or immediate clarity. For example, although most participants reported at least one illustration as confusing (Figure 4C), the image most frequently identified (Figure 3E, depiction of genetic variation across Indigenous populations globally) was intentionally retained following participant feedback emphasizing the importance of understanding how local communities fit within a broader Indigenous context, although our study cannot fully disentangle whether this confusion arose from the visual representation itself or from the underlying concept being communicated.

Second, we found that participants strongly preferred illustrations that reflected familiar people, environments, and lived experiences. During early phases of development, both Turkana and Orang Asli participants expressed dissatisfaction with illustrations that used generic human figures or attempted to represent diversity through a range of skin tones; instead, participants wanted to see individuals who looked like themselves and contexts that reflected their own communities. This finding aligns with prior work demonstrating that analogies grounded in shared experiences facilitate comprehension of abstract biological concepts. For example, digital storytelling in Alaska Native communities has been shown to be a culturally respectful and engaging approach to science communication, particularly when narratives are rooted in local knowledge and experience [55]. More broadly, this finding mirrors patterns observed in ethical governance frameworks: while global principles such as the United Nations Declaration on the Rights of Indigenous Peoples (UNDRIP) and the CARE principles [56,57] provide important guidance, their effective application requires attention to the specific histories, cultures, and priorities of individual Indigenous communities, which are not homogeneous [23,58,59].

Finally, the heterogeneity we observed in engagement with and understanding of the illustrations further reinforces the need to move beyond a generalizable, “one size fits all” approach. Although participants overall showed significant interest in learning about genetics, strong engagement, and perceived gains in understanding, our quantitative analyses revealed modest but consistent association between responses and urbanicity, education, age, and sex. Education level in particular emerged as a key predictor, likely reflecting differential exposure to genetics-related topics. Individuals with more formal education were more likely to report that they could explain the illustrations to a friend, potentially reflecting greater prior familiarity, but also that they would like to learn more about these topics, suggesting that prior educational exposure may also foster greater confidence and interest in scientific information. Additionally, individuals in less urbanized locations were more likely to report that the illustrations helped them understand why researchers wanted to study DNA, consistent with this being an early exposure to these concepts. Furthermore, we observed sex-based differences in engagement and interpretation, suggesting that learning preferences and perceived relevance may vary across social roles and experiences, a phenomenon identified in previous literature [60,61].

Our study has several limitations. First, while illustrations were piloted with both Turkana and Orang Asli communities, our decision to switch to community-specific illustrations resulted in formal evaluations only being conducted with Orang Asli participants, making us unable to compare illustration effectiveness across populations. Second, interviews relied on retrospective self-assessment rather than objective baseline measures, which may introduce recall bias [62]. Finally, because the final version of illustrations were delivered as live presentations by OA HeLP research assistants, slight variation in phrasing across presentations may have influenced responses. However, we view this flexibility as a strength rather than a weakness, reflecting real-world conditions in which engagement is relational, interactive, and adaptive, rather than standardized. Additionally, because illustrations were presented alongside verbal explanations, we are unable to determine the extent to which visual materials themselves contributed to comprehension relative to verbal explanations alone. More broadly, visual limitations themselves have inherent limitations. For example, several community members expressed interest in animated versions of the illustrations, suggesting that motion and narration may further enhance clarity for complex biological processes. While animation would allow for more dynamic explanations, producing high-quality animated materials requires substantial financial resources, technical expertise, and reliable technological infrastructure. Moreover, engagement formats must remain concise; in our experience, community members are unlikely to engage with materials longer than 10-15 minutes. These considerations highlight the balance researchers must strike between ideal communication tools and practical constraints of time, funding, and local context. Rather than serving as comprehensive explanations of research in themselves, illustrations are best understood as accessible entry points that can prompt further discussion and ongoing dialogue.

Despite these limitations, we hope that reporting on the challenges and procedures involved in this work offers guidance for others. We have identified three practical implications, which largely echo prior themes [19,20,30]. First, effective communication materials should be treated as evolving resources rather than finalized products, with time and resources allocated for iterative revision. We acknowledge (and experienced) that this can be challenging given the difficulty of securing dedicated resources (e.g., grant funding) for such work. Second, community-specific tailoring should be considered a critical aspect to illustration design. Third, it is important to evaluate engagement materials not only for comprehension, but also for whether they resonate with participants’ interests, values, and goals for engaging with researchers. We have already begun applying these principles to other OA HeLP and THGP initiatives, including recent results-return efforts. An additional area for future work is to examine whether participatory communication exercises such as this influence broader trust in science and researchers, particularly among individuals who may have had negative prior experiences with health projects or biological sample collection. As genomic research continues to occur alongside historically underrepresented, Indigenous populations, approaches like these aim to offer a pathway for building trust and fostering mutual understanding.

## Data availability statement

OA HeLP’s highest priority is the minimization of risk to study participants. OA HeLP adheres to the ‘CARE Principles for Indigenous Data Governance’ (Collective Benefit, Authority to Control, Responsibility, and Ethics). OA HeLP is also committed to the ‘FAIR Guiding Principles for scientific data management and stewardship’ (Findable, Accessible, Interoperable, Reusable). To adhere to these principles while minimizing risks, individual-level data are stored in the OA HeLP protected data repository, and are available through restricted access. Requests for de-identified, individual-level data should take the form of an application that details the exact uses of the data and the research questions to be addressed, procedures that will be employed for data security and individual privacy, potential benefits to the study communities and procedures for assessing and minimizing stigmatizing interpretations of the research results. Requests for de-identified, individual-level data will require institutional IRB approval (even if exempt). OA HeLP is committed to open science and the project leadership is available to assist interested investigators in preparing data access requests (see orangaslihealth.org for further details and contact information).

Summaries of all presented data are in the Supplementary Materials. Scripts used for these analyses can be found on GitHub (https://github.com/audreyarner/genetic_illustrations).

## Acknowledgements

Above all else, we thank the Orang Asli and Turkana participants who have generously allowed us to work in their communities, as well as for their hospitality and support of this project. We also thank members of the Orang Asli Health and Lifeways Project and Turkana Health and Genomics Project who reviewed the early versions of the illustrations. We are also grateful to Jada Benn Torres and the members of the Lea Lab for their feedback and support.

