## Supplementary Information for "Development and assessment of tailored illustrations to enhance community understandings of genetics topics"

This supplement consists of:

**Supplementary Methods**

**Supplementary Information**
- Supplementary Information 1. English Interview

**Supplementary Text**

**Supplementary Figures 1-7**
- Supplementary Figure 1. Thematic map for the information participants thought was in their blood prior to seeing the illustrations
- Supplementary Figure 2. Thematic map for responses about what each participant learned.
- Supplementary Figure 3. Thematic map for responses about why a chosen image was the participant’s favorite.
- Supplementary Figure 4. MCA scree plot

**Supplementary Tables**
- Supplementary Table 1. Description of each illustrated concept
- Supplementary Table 2. Demographic information for participants taking final interview
- Supplementary Table 3. Percent interrater agreement for codes
- Supplementary Table 4. Genetics-related topics participants expressed further interest in
- Supplementary Table 5. Binomial model results for question response ~ 1
- Supplementary Table 6. MCA dimension model results as a function of sex, age, highest education level, and urbanicity score
- Supplementary Table 7. Question response model results as a function of sex, age, highest education level, and urbanicity score
- Supplementary Table 8: Question response model results as a function of sex, age, binary of any formal education, and urbanicity score

Supplementary Information 1: English Interview

**Part 1: Demographic information**

1. Date
2. Interviewer
3. Local name
4. Name on IC card
5. Sex
6. Can I take a picture?
7. Do you know how old you are?
    a. If yes, age
    b. If no, estimated age
8. Village
9. How many years of schooling have you had?
10. Highest level of schooling *(none, primary, secondary, college)*

**Part 2: Prior knowledge**

11. Would you like to know more about blood?
12. Do you think there is information about health in your blood?
    a. If yes, what type of information did you think was in your blood prior to seeing the images?

**Part 3: Thoughts after viewing the illustrations:**

13. What image was your favorite?
14. Why was that image your favorite?
15. Were there images you did not understand?
    a. If yes, which ones?
16. If you could learn more about an image, which would it be?
17. What else would you like to know about your blood? *(health & disease/relatedness/specifics about DNA/similarity and differences to other populations/other)*
18. What is one thing you learned from the illustrations?
19. Do you think you could explain at least one of these illustrations to a friend?
20. I think I know more about DNA now than I did before seeing the illustrations

21. The presentation and illustrations helped me understand why a scientist would want to study blood
22. The images helped me understand why researchers want to study Orang Asli
23. The illustrations were hard to understand
24. I would look at the illustrations again
25. I would tell a friend to look at the illustrations
26. I would like to learn more about what I learned today

### Supplementary Text

#### *Urbanicity score generation*

Orang Asli live across a wide lifestyle gradient, which we have previously described using a location-based "urbanicity" score [1,2]. Using a location-level scale is advantageous because it gives a representation of the resources available across the community (e.g., an individual may not own a television but may have access through families or friends). To quantify individuals' exposure to urban centers and urban infrastructure, we used a scale first proposed by Novak et al [3]. This scale was tested in Orang Asli and predicted cardiometabolic health better than other measures of urbanicity. Scale construction is shown below and population density was estimated from NASA's Gridded Population of the World resource with a resolution of 2.5 arc-minutes [4]. The urbanicity scores values ranged from 13.802-24.636, modeled as a continuous variable in linear models during our analysis.

| **Population density (estimated number of people per square kilometer)** | **Contribution to scale** |
|---|---|
| 0-100 | 1 |
| 100-200 | 2 |
| 200-300 | 3 |
| 300-400 | 4 |
| 400-500 | 5 |
| 500-1000 | 6 |
| 1000-2000 | 7 |
| 2000-3000 | 8 |
| 3000-4000 | 9 |
| 4000-6000 | 10 |
| 6000-8000 | 11 |
| 8000-10000 | 12 |
| 10000-15000 | 13 |
| 15000-20000 | 14 |
| >20000 | 15 |
| **Occupation** | |
| Proportion of the population involved in non-wage labor | 10 - (10 x (proportion)) |
| **Built Environment** | |
| Proportion of households with flush toilets | 5 x proportion |

| | |
|---|---|
| Proportion of households with electricity | 5 x proportion |
| **Communication/market-derived items** | |
| Proportion of households with mobile phone | 5 x proportion |
| Proportion of households with television | 5 x proportion |
| **Education** | |
| Proportion of surveyed individuals > 40 years with any level of formal education | 10 x proportion |
| Proportion of surveyed individuals < 40 years with any level of formal education | 10 x proportion |

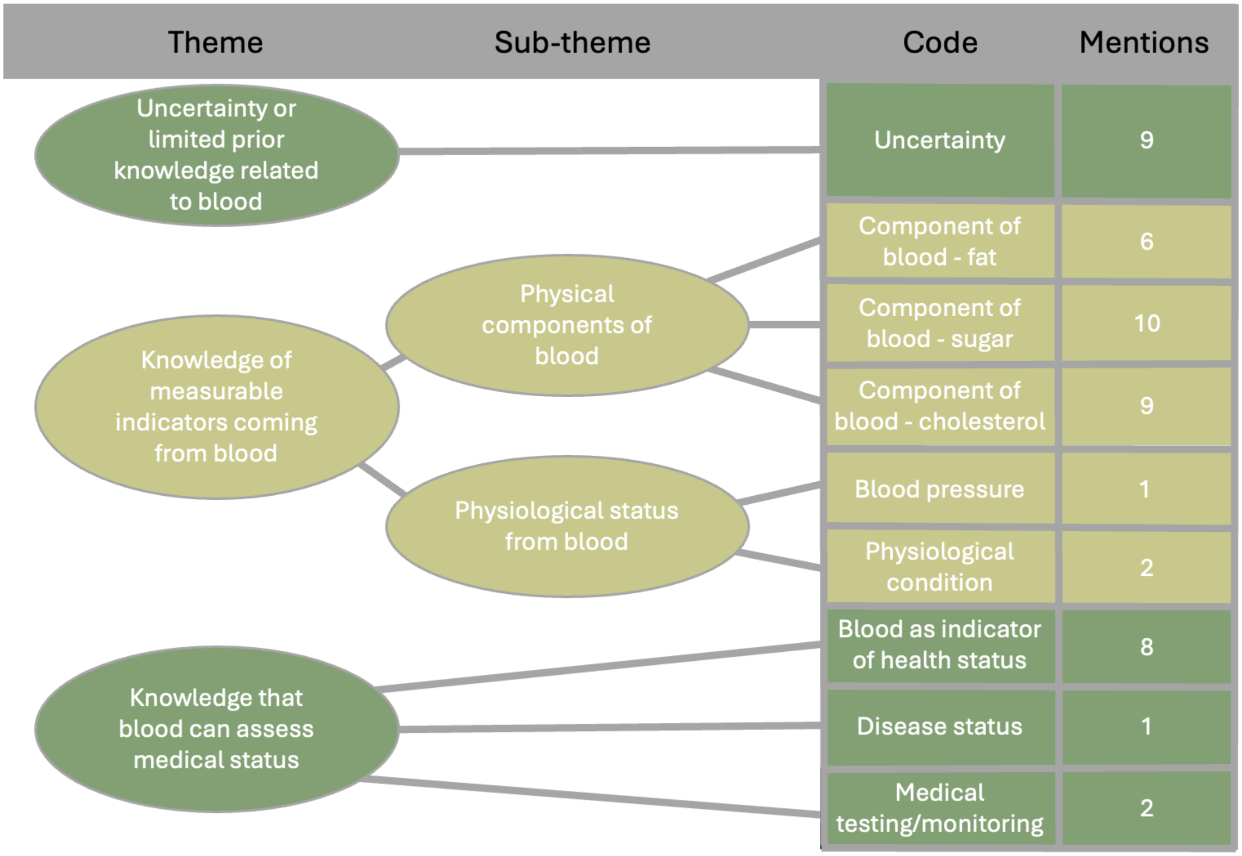


**SI Figure 1. Thematic map for the information participants thought was in their blood prior to seeing the illustrations.** Each theme and sub-theme are listed, as well as the codes associated with each. Mentions refers to the total number of times a code was recorded across all individuals.

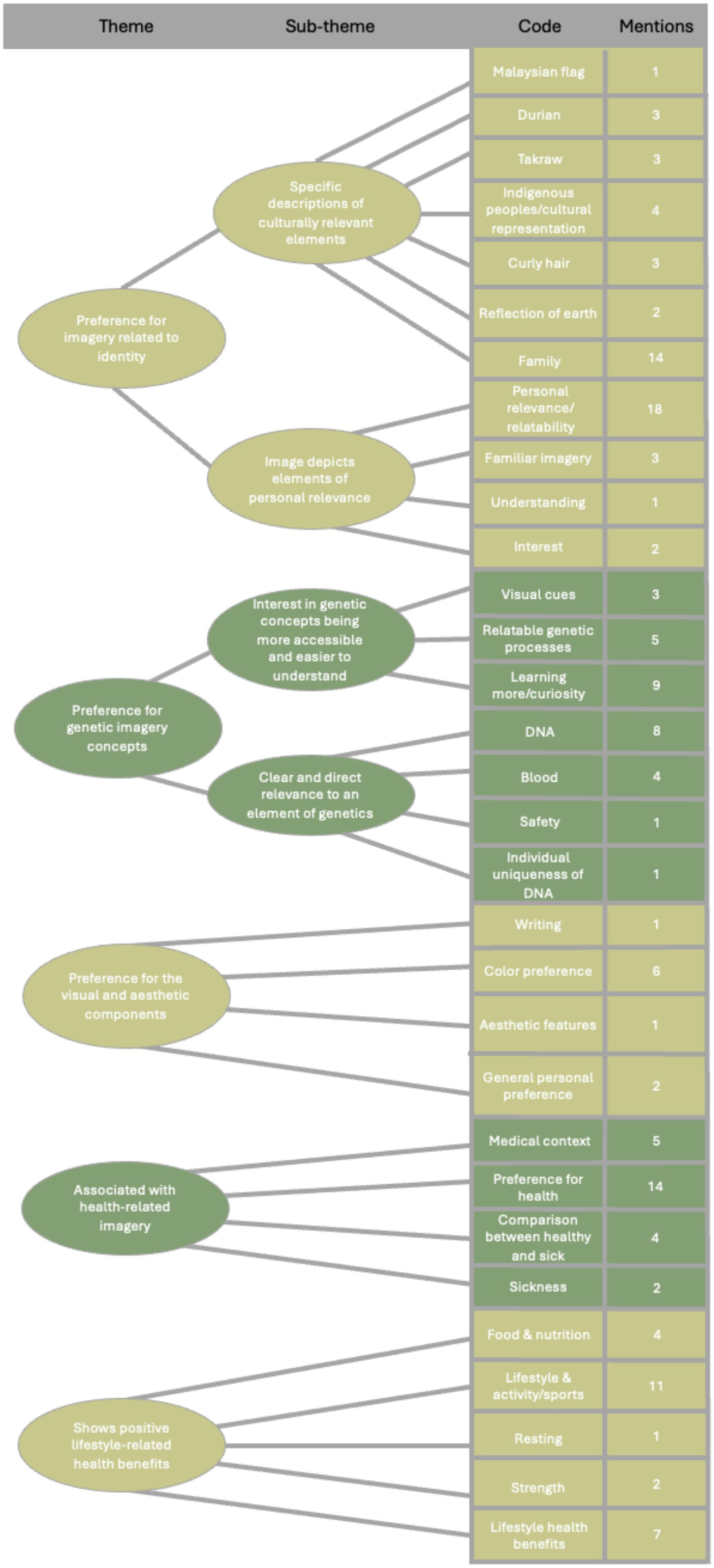


**SI Figure 2. Thematic map for responses about what each participant learned.** Each theme and sub-theme are listed, as well as the codes associated with each. Mentions refers to the total number of times a code was recorded across all individuals.

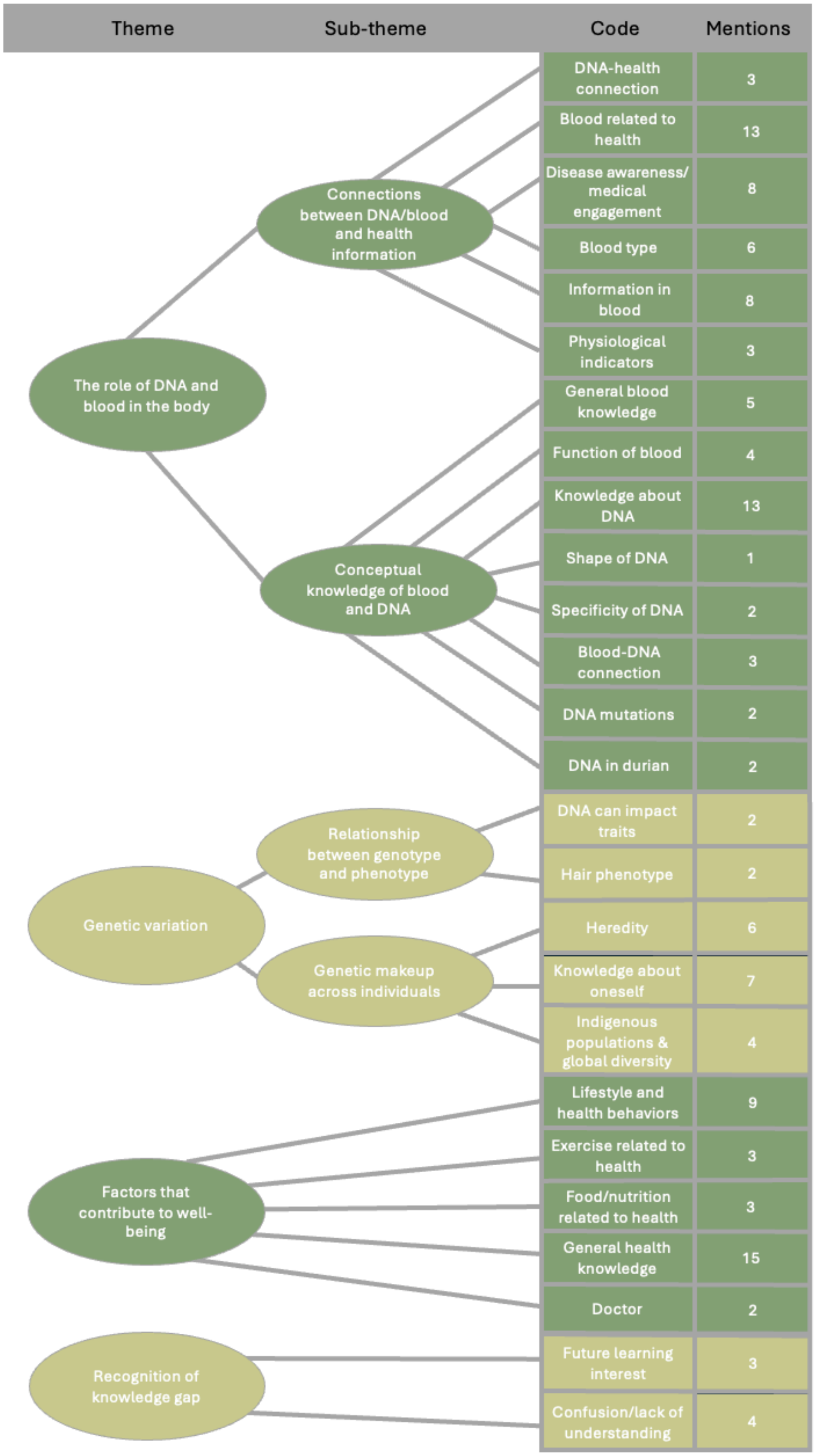


**SI Figure 3. Thematic map for responses about why a chosen image was the participant's favorite.** Each theme and sub-theme are listed, as well as the codes associated with each. Mentions refers to the total number of times a code was recorded across all individuals.

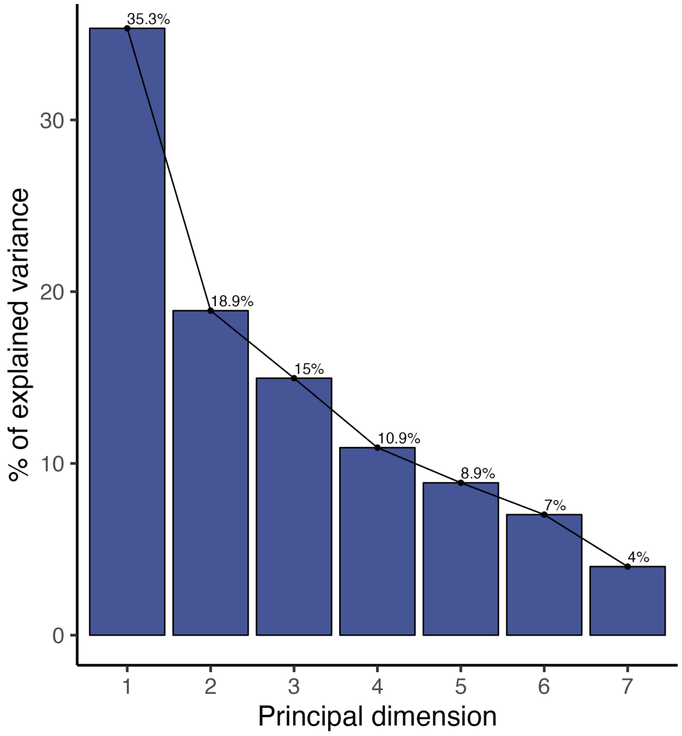


**SI Figure 3. MCA scree plot.** Scree plot showing the proportion of variance explained by the top six dimensions of the multiple correspondence analysis (MCA).

SI Table 1: Description of each illustrated concept

| Question | Concept illustrated | Example of Orang Asli imagery used |
|---|---|---|
| What is DNA? | DNA carries information inherited from parents and shared across living things | Hair texture variation among family members, durian varieties |
| Can DNA affect your health? | Some DNA changes have effects on health while most do not | Traditional clothing, housing, and sleep mat |
| What can scientists learn from DNA? | How changes in DNA relate to health and ancestry | Picture of Orang Asli along with other Indigenous populations |
| Besides DNA, what else do you study in my blood? | Blood contains measurable components like sugars and fats | Nasi lemak (traditional dish) with less sugars than a burger in the city |
| Why are scientists interested in markers of health? | Links between lifestyle change and biomarkers | Physical activity of an individual playing takraw (traditional ball game) |
| Who has access to my DNA? | Data access and research ethics | Include Kuala Lumpur (capital city of Malaysia) in the background |

SI Table 2: Demographic information for participants taking final interview

| Location | # participants | Ethnolinguistic group | Sex | | Urbanicity score | Highest level of formal education | | | |
|---|---|---|---|---|---|---|---|---|---|
| | | | # F | # M | | None | Primary school | Secondary school | University |
| A | 21 | Temiar | 8 | 13 | 19.156 | 0 | 11 | 9 | 1 |
| B | 10 | Semai | 8 | 2 | 24.691 | 0 | 3 | 7 | 0 |
| C | 17 | Semai | 13 | 4 | 24.636 | 12 | 3 | 2 | 0 |
| D | 20 | Semai | 11 | 9 | 13.802 | 12 | 3 | 5 | 0 |
| E | 8 | Temiar | 8 | 0 | 21.920 | 1 | 2 | 5 | 0 |
| F | 16 | Temiar | 5 | 11 | 19.645 | 6 | 5 | 5 | 0 |

SI Table 3: Percent interrater agreement for codes

| Question | Percent interrater agreement |
|---|---|
| What type of information did you think was in your blood prior to seeing the images? | 100% |
| Why was that image your favorite? | 85.7% |
| What is one thing you learned after seeing the illustrations? | 86.2% |

SI Table 4: Genetics-related topics participants expressed further interest in

| Topic | Individuals |
|---|---|
| Health and disease | 45 |
| Relatedness | 42 |
| Specifics about DNA | 17 |
| Similarities and differences to other populations | 26 |
| Other | 13 |

SI Table 5: Binomial model results for question response ~ 1

| Question | P-value | FDR |
|---|---|---|
| 1. I think I know more about DNA now | $7.48 \times 10^{-8}$ | $1.2 \times 10^{-7}$ |
| 2. The illustrations helped me understand more about genetics | $6.82 \times 10^{-10}$ | $2.73 \times 10^{-9}$ |
| 3. The illustrations helped me understand why researchers want to study DNA | $1.47 \times 10^{-8}$ | $2.94 \times 10^{-8}$ |
| 4. I could explain the illustrations to a friend | $5.45 \times 10^{-4}$ | $6.23 \times 10^{-4}$ |
| 5. I would look at the illustrations again | $1.18 \times 10^{-9}$ | $3.15 \times 10^{-9}$ |
| 6. I would tell a friend to look at the illustrations | $1.04 \times 10^{-7}$ | $1.38 \times 10^{-7}$ |
| 7. I would like to learn more about genetics | $5.18 \times 10^{-10}$ | $2.73 \times 10^{-9}$ |
| 8. The illustrations were hard to understand | $4.08 \times 10^{-3}$ | $4.08 \times 10^{-3}$ |

SI Table 6: MCA dimension model results as a function of sex, age, highest education level, and urbanicity score

| Outcome | Covariate | Beta | SE | P-value | FDR |
|---|---|---|---|---|---|
| Dimension 1 | sex | -0.1299 | 0.1291 | 0.3172 | 0.3966 |
| Dimension 1 | age | -0.0003 | 0.0058 | 0.9575 | 0.9575 |
| Dimension 1 | highest education level | -0.1654 | 0.0897 | 0.0686 | 0.1267 |
| Dimension 1 | urbanicity score | -0.0287 | 0.0159 | 0.0760 | 0.1267 |
| Dimension 2 | sex | -0.1279 | 0.0954 | 0.1835 | 0.4083 |
| Dimension 2 | age | -0.0005 | 0.0043 | 0.9024 | 0.9024 |
| Dimension 2 | highest education level | -0.0407 | 0.0662 | 0.5402 | 0.6753 |
| Dimension 2 | urbanicity score | 0.0243 | 0.0118 | 0.0422 | 0.2113 |

SI Table 7: Question response model results as a function of sex, age, highest education level, and urbanicity score

| question | variable | beta | st_error | p_value | fdr |
|---|---|---|---|---|---|
| 1 | Age | 0.003215198846 | 0.003987596307 | 0.4222684597 | 0.6353954644 |
| 1 | Sex | 0.08775992322 | 0.08678117776 | 0.3146875469 | 0.5300000789 |
| 1 | Education level | 0.07709321036 | 0.06114181451 | 0.2107187443 | 0.4495333212 |
| 1 | Urbanicity score | 0.01239752744 | 0.01101922991 | 0.2636490189 | 0.4962805062 |
| 2 | Age | 0.002569444924 | 0.003289130728 | 0.4368343817 | 0.6353954644 |
| 2 | Sex | 0.01726795776 | 0.07153068339 | 0.8098148831 | 0.9886435192 |
| 2 | Education level | 0.1035889945 | 0.05019269335 | 0.04204836923 | 0.2242579692 |
| 2 | Urbanicity score | 0.000779754015 | 0.009008618591 | 0.9312252974 | 0.9886435192 |
| 3 | Age | -0.00378675432 | 0.003637300459 | 0.3007186767 | 0.5300000789 |
| 3 | Sex | 0.1212770542 | 0.0791577666 | 0.1291279073 | 0.3779761548 |
| 3 | Education level | 0.01146336813 | 0.05577072825 | 0.8376268989 | 0.9886435192 |
| 3 | Urbanicity score | -0.02508831072 | 0.0100512306 | 0.01444456799 | 0.1155565439 |
| 5 | Age | -0.00381125303 | 0.003357394538 | 0.2594530834 | 0.4962805062 |
| 5 | Sex | 0.1167561207 | 0.07365913728 | 0.116617101 | 0.3779761548 |
| 5 | Education level | -0.00659074337 | 0.05160876738 | 0.8986798777 | 0.9886435192 |
| 5 | Urbanicity score | 0.01534391757 | 0.009274073925 | 0.1016722227 | 0.3779761548 |
| 4 | Age | -0.00243603666 | 0.004255429826 | 0.568491036 | 0.7276685261 |
| 4 | Sex | 0.1294953631 | 0.09260997949 | 0.1655811927 | 0.4075844742 |
| 4 | Education level | 0.1724907029 | 0.06524850588 | 0.009729209266 | 0.1155565439 |
| 4 | Urbanicity score | 0.02571077747 | 0.01175935476 | 0.03147017765 | 0.201409137 |
| 6 | Age | -0.00005680097 | 0.003979006807 | 0.9886435192 | 0.9886435192 |
| 6 | Sex | -0.00259617953 | 0.08653382906 | 0.9761350516 | 0.9886435192 |
| 6 | Education level | 0.09284262475 | 0.06072031946 | 0.1299293032 | 0.3779761548 |
| 6 | Urbanicity score | 0.01609724835 | 0.01089812405 | 0.1433106621 | 0.3821617655 |
| 7 | Age | 0.00242771227 | 0.003097547943 | 0.4353136798 | 0.6353954644 |
| 7 | Sex | 0.09126802697 | 0.0674112518 | 0.1792752397 | 0.4097719764 |
| 7 | Education level | 0.1225525723 | 0.04749470288 | 0.01154690668 | 0.1155565439 |
| 7 | Urbanicity score | 0.01630061276 | 0.008559691184 | 0.06016993016 | 0.2750625379 |
| 8 | Age | 0.000324793532 | 0.004630859905 | 0.9442457705 | 0.9886435192 |
| 8 | Sex | 0.281308985 | 0.1007803814 | 0.006451787694 | 0.1155565439 |
| 8 | Education level | -0.04138916358 | 0.07100497531 | 0.5614650503 | 0.7276685261 |
| 8 | Urbanicity score | -0.00854889984 | 0.01279680942 | 0.5058705599 | 0.7038199094 |

SI Table 8: Question response model results as a function of sex, age, binary of any formal education, and urbanicity score

| question | variable | beta | st_error | p_value | fdr |
|---|---|---|---|---|---|
| 1 | Age | 0.02995350554 | 0.02714221457 | 0.2697769637 | 0.6640663723 |
| 1 | Sex | 0.3821043193 | 0.5940945987 | 0.5201132552 | 0.7236358334 |
| 1 | Any formal education | 1.230417692 | 0.7253674397 | 0.08983510753 | 0.3699191318 |
| 1 | Urbanicity score | 0.07313271523 | 0.06931281806 | 0.291374572 | 0.6659990216 |
| 2 | Age | 0.02628542464 | 0.03105149423 | 0.3972678621 | 0.669082715 |
| 2 | Sex | -0.1217648177 | 0.7410769965 | 0.8694887494 | 0.9970277789 |
| 2 | Any formal education | 1.857555014 | 0.8735663488 | 0.03346963163 | 0.2142056424 |
| 2 | Urbanicity score | 0.003155554442 | 0.08317771413 | 0.9697375147 | 0.9970277789 |
| 3 | Age | -0.02591328983 | 0.02678376965 | 0.3332942776 | 0.6665885552 |
| 3 | Sex | 1.039295912 | 0.7362200958 | 0.1580487154 | 0.438810455 |
| 3 | Any formal education | 0.01760004629 | 0.7384813043 | 0.9809860103 | 0.9970277789 |
| 3 | Urbanicity score | -0.186043529 | 0.08724501274 | 0.03297184619 | 0.2142056424 |
| 5 | Age | -0.03006894207 | 0.03026122588 | 0.3203953049 | 0.6665885552 |
| 5 | Sex | 1.124136685 | 0.7670748382 | 0.1427886364 | 0.438810455 |
| 5 | Any formal education | 0.003194556715 | 0.8575679715 | 0.9970277789 | 0.9970277789 |
| 5 | Urbanicity score | 0.1185098531 | 0.07793200425 | 0.1283394742 | 0.438810455 |
| 4 | Age | -0.00210126420[illegible] | 0.02406540105 | 0.9304213389 | 0.9970277789 |
| 4 | Sex | 0.4151990803 | 0.5686158728 | 0.4652724891 | 0.6794378085 |
| 4 | Any formal education | 1.96738656 | 0.6662191963 | 0.003146388108 | 0.1006844195 |
| 4 | Urbanicity score | 0.1253083181 | 0.06644466295 | 0.05930776574 | 0.316308084 |
| 6 | Age | -0.00762875365[illegible] | 0.02568582798 | 0.7664646442 | 0.9810747445 |
| 6 | Sex | -0.09279852123 | 0.5796457386 | 0.872806054 | 0.9970277789 |
| 6 | Any formal education | 0.6426137546 | 0.7230291326 | 0.3741213845 | 0.669082715 |
| 6 | Urbanicity score | 0.09505917563 | 0.06839195371 | 0.1645539206 | 0.438810455 |
| 7 | Age | 0.02957446935 | 0.03363283244 | 0.3792205885 | 0.669082715 |
| 7 | Sex | 0.6346633354 | 0.79364248 | 0.4238938013 | 0.6782300821 |
| 7 | Any formal education | 2.202212122 | 0.9479764787 | 0.02017559607 | 0.2142056424 |
| 7 | Urbanicity score | 0.1435125227 | 0.08529924642 | 0.09247978295 | 0.3699191318 |
| 8 | Age | 0.009319825233 | 0.02227953018 | 0.6757180196 | 0.9009573594 |
| 8 | Sex | 1.353043418 | 0.5254514954 | 0.01002368783 | 0.1603790052 |
| 8 | Any formal education | -0.04769608231 | 0.6166097162 | 0.9383434118 | 0.9970277789 |
| 8 | Urbanicity score | -0.04424962242 | 0.06085068835 | 0.4671134933 | 0.6794378085 |
